# Brain Network Segregation is Associated with Drug Use Severity in Individuals with Opioid Use Disorder

**DOI:** 10.1101/2025.04.02.646889

**Authors:** Nathan M. Hager, Xinying Wang, Astrid P. Ramos-Rolón, Anna Rose Childress, Daniel D. Langleben, Corinde E. Wiers, Zhenhao Shi

## Abstract

Opioid use disorder (OUD) is associated with altered brain network connectivity, particularly in the fronto-parietal (FPN), default mode, and salience (SN) networks. At rest, brain networks that are distinct from each other but are partially connected can optimize neural efficiency and support cognitive performance. Previous research found lower network segregation in people with cognitive impairment, alcohol use disorder, and as people age. Here, we examined “brain network segregation”—a graph theory-based metric of the network integration/segregation balance—in individuals with OUD and hypothesized that recent drug use severity would be linked to reduced network segregation. Forty adults with OUD completed resting-state functional magnetic resonance imaging, the drug use severity subscale of the Addiction Severity Index, and measures of cognition (IQ and working memory), mood, and affect. We grouped 264 brain regions into 10 networks, categorized as “association” (higher-order cognition) or “sensorimotor” (sensory and motor) networks. Regression analyses showed that drug use severity predicted lower brain network segregation in the association networks, with the FPN and SN driving this effect. Age predicted lower brain network segregation in the sensorimotor networks, while an interaction with age showed that drug use severity only predicted lower sensorimotor network segregation in younger adults. Cognition did not relate to brain network segregation, but positive affect related to greater SN segregation. Brain network segregation remained stable across OUD treatment. These findings elucidate alterations in brain network segregation related to drug use severity in people with OUD, which may contribute to cognitive impairment and accelerated brain aging.

## Introduction

In recent years, approximately six million people in the United States experienced past-year opioid use disorder (OUD) (1,2). Risks of OUD include opioid overdoses and associated medical complications and mortality, social dysfunction, cognitive deficits, and poor mental health (3–6). Underlying these poor outcomes, chronic substance use affects neural functioning in a variety of domains—including reward, emotion, stress, sensorimotor, learning, and executive function (7). Despite the multifaceted effects of substance use, effective treatments for OUD mostly focus on reducing physical withdrawal (8) and face challenges with adherence and retention (9).

Our understanding of the widespread brain effects of OUD is critical to expanding therapeutic targets for this disorder (10). Previous work has shown that resting-state functional connectivity (rsFC) assessed by functional magnetic resonance imaging (fMRI) can shed light on the impact of substance use disorders (SUDs) on the configuration of and interplay between distinct brain networks (i.e., widespread brain areas defined by shared functional domains) (11). People with SUDs have alterations within and between a variety of neural networks including the fronto-parietal network (FPN), default mode network (DMN), salience network (SN), affective/reward network, and somatosensory network (12). The most common networks implicated in OUD are the same as those most implicated in SUDs broadly: FPN, DMN, and SN (13–16). Analysis of rsFC network may help parse subgroups of SUD (17), identify which networks protect from or contribute to SUD (18), and improve treatment-matching and intervention targets (19).

Researchers have applied rsFC to compute graph theory-based metrics of global brain network organization, including modularity and clustering coefficient. Such measures capture the overall topological architecture and efficiency of brain networks (20). Brain network segregation (also known as system segregation) is a metric that quantifies the balance between having segregated and integrated brain networks (i.e., dense within-network connections and sparser between-network connections) (21). This property of distinct yet partially integrated neural networks is a fundamental aspect of the human brain that balances network specialization with global integration (22). Wig (2017) proposed that this balance of network segregation is beneficial because it preserves functional heterogeneity/specialization in the brain, contributes to energetically efficient neural adaptation, and promotes resilience to neural damage. Chan and colleagues (2014) showed that brain network segregation predicts aging of the human brain better than other network metrics. To gain some specificity of where network segregation differences appear, previous studies have found meaningful differences when separately examining the “association” and “sensorimotor” networks (21,24). Association networks are involved in higher-order cognition and information integration while sensorimotor networks are involved in processing incoming sensory and outgoing motor information.

Studies consistently show that resting-state brain networks become less segregated over the course of normal adult aging (21,24), and less segregation is often associated with poorer cognitive performance (21,24–26). Lower network segregation has also been associated with severity of dementia (27). Only one study has examined this network segregation metric in SUD. As expected, individuals with alcohol use disorder had lower brain network segregation in the association and sensorimotor networks compared to healthy control participants (28). Studies using connectivity metrics similar to network segregation (e.g., small-worldness, clustering coefficient) have shown similar findings, such that SUDs may be characterized by less efficient and hyperconnected networks (29,30). No previous research has examined network segregation in OUD, but one study found that people with OUD have decreased small-worldness and clustering coefficient (31). These findings indicate that OUD is characterized by more random brain organization, suggesting a possible reduction in network segregation. Examining network segregation in people with OUD may provide insight into the effects of chronic opioid use on widespread brain network organization, efficiency, and adaptability.

We present analyses of brain network segregation in relation to drug use severity in a group of individuals with OUD. These are secondary analyses of data from a clinical trial of injectable, once-monthly extended-release naltrexone (XR-NTX) for the treatment of OUD (32). By testing network segregation against a continuous measure of drug use severity, we sought to understand the sensitivity of network segregation to OUD symptomatology. We hypothesized that drug use severity and age would independently predict lower brain network segregation in OUD, particularly in the FPN, DMN, and SN as these networks are consistently altered in OUD. Thus, we extend previous research by examining network segregation in OUD and testing the effects both broadly and within specific networks. In exploratory analyses, we expected that network segregation would increase across XR-NTX treatment and drug use severity would decrease. We further explored the associations of brain network segregation with emotion (depression, anxiety, and negative affect, and positive affect) and cognition (IQ and working memory).

## Method

### Participants

Treatment-seeking individuals with OUD from the greater Philadelphia area volunteered to receive up to three monthly XR-NTX injections. Data were collected between 2012 and 2014. Of the initial 46 participants, we obtained the final sample of *N* = 40 after excluding for incomplete scans (*n* = 4), excessive head motion (*n* = 1), and incomplete Addiction Severity Index (*n* =1). See Table 1 for demographics and baseline characteristics.

**Table 1.** Participant Demographics and Mean Scores (N = 40)

|  | *n* | Range | Mean (*SD*) |
| --- | --- | --- | --- |
| Age (years) | 40 | 19 – 56 | 29.33 (9.21) |
| Education (years) | 40 | 7 – 19 | 14.00 (2.18) |
| Drug severity | 40 | 0.08 – 0.51 | 0.29 (0.11) |
| Sensorimotor network segregation | 40 | 0.29 – 0.60 | 0.46 (0.08) |
| Association network segregation | 40 | 0.18 – 0.45 | 0.34 (0.06) |
| Depression<sup>1</sup> | 39 | 0 – 19 | 6.97 (5.01) |
| Anxiety<sup>2</sup> | 38 | 0 – 20 | 6.76 (5.32) |
| Positive affect<sup>3</sup> | 24 | 12 – 49 | 29.29 (9.01) |
| Negative affect<sup>3</sup> | 24 | 12 – 44 | 25.75 (9.02) |
| IQ<sup>4</sup> | 40 | 80 – 130 | 102.43 (12.20) |
| Working memory<sup>5</sup> | 26 | 75 – 117 | 100.12 (11.45) |
|  |  | % |  |
| Sex |  |  |  |
| Female | 11 | 27.5% |  |
| Male | 29 | 72.5% |  |
| Race |  |  |  |
| Asian | 1 | 2.5% |  |
| Black | 3 | 7.5% |  |
| Hispanic | 3 | 7.5% |  |
| White | 33 | 82.5% |  |

*Note:* ASI = Addiction Severity Index (Drug Composite Score); HAM-D = Hamilton Depression Rating Scale; HAM-A = Hamilton Anxiety Rating Scale, PANAS = Positive Affect Negative Affect Schedule.

### Study Procedures

This study was approved by the university’s Institutional Review Board, and all participants signed voluntary informed consent. Prior to their baseline MRI scan, participants underwent outpatient opioid detoxification. Following the baseline scan (T1), the on-treatment MRI scan (T2) and post-treatment MRI scan were completed a mean of 13.0 (*SD* = 7.7) and 112.2 (*SD* = 21.4) days later, respectively. The first XR-NXT injection occurred after the baseline scan (see Supplemental Materials).

## Measures

### Drug Use Severity

The Addiction Severity Index-5^th^ Edition (33) and its Drug Composite score (ASI-drug) was used to measure drug use severity. The ASI-drug focuses on the past 30 days and all abused substances except alcohol. It is comprised of self-reported days of use, number of drug problems, and ratings for problem severity and treatment need. Higher scores indicate higher recent drug use severity.

#### Depression and Anxiety

Depression severity was assessed by the Hamilton Depression Rating Scale (34) which is a 17-item clinician-rated measure. The internal consistency, assessed via Cronbach’s alpha (α) was acceptable in the current sample (α = .73). We assessed anxiety with the 14-item, clinician-rated Hamilton Anxiety Rating Scale (35). Internal consistency was good in the current sample (α = .80). Higher scores indicated greater symptom severity.

#### Positive and Negative Affect

Positive and negative affect was assessed by the Positive Affect Negative Affect Schedule (PANAS) (36). The PANAS contains two 10-item subscales to measure positive and negative affect separately. Participants self-reported the frequency that they have experienced various positive (e.g., interested, proud) and negative (e.g., afraid, irritable) emotions. Internal consistency was good for both the positive (α = .89) and negative (α = .89) subscales.

#### Cognition

General cognitive function (IQ) was calculated from the Vocabulary and Matrix Reasoning subtests of the Wechsler Abbreviate Scale of Intelligence-1^st^ edition (WASI) (37). We measured working memory using the Working Memory Index (WMI) of the Wechsler Assessment of Intelligence Scale-III (38). IQ and WMI scores were age-adjusted and standardized to the norming sample.

### MRI Acquisition and Preprocessing

MRI scans were collected on a Siemens Trio 3T system (Siemens AG, Erlangen, Germany). The rs-fMRI data were collected using a whole-brain, single-shot gradient-echo echo-planar sequence with repetition time (TR)/echo time (TE) = 2000/30 ms, field of view (FOV) = 220×220 mm^2^, matrix = 64×64, slice thickness/gap = 4.5/0 mm, 32 slices, effective voxel resolution of 3.4×3.4×4.5 mm^3^, flip angle (FA) = 90°. For structural imaging, the magnetization-prepared rapid acquisition gradient echo sequence acquired high-resolution T1-weighted whole-brain images with TR/TE = 1510/3.71 ms, FOV = 256×192 mm^2^, matrix = 256×192, slice thickness/gap = 1/0 mm, 160 slices, effective voxel resolution of 1×1×1 mm^3^, FA = 9°. The rs-fMRI data were preprocessed in MATLAB using a pipeline adapted from Ciric and collegues (2018). This consisted of removing the first five scans, estimation of the 24 motion parameters (including the six raw motion parameters, six framewise displacement [FD] parameters, the square of the raw motion parameters, and the square of the [FD] parameters), identification of FD timepoints (> 0.5 mm), identification slice time correction, motion correction, coregistration and segmentation of the structural images, skull stripping, computation of DVARS and identification of DVARS outliers using the procedure described in Afyouni & Nichols (2018), despiking using AFNI’s 3dDespike, removal of polynomial trends (order=3), extraction of nuisance signals from the voxels located within the top 10% of the deepest tissue of the white matter and the cerebrospinal fluid, interpolation of FD and DVARS outlier timepoints using Lomb-Scargle periodogram, bandpass filtering at 0.01–0.1 Hz of the images and covariates (i.e., 24 motion parameters and two nuisance signals), regressing out the filtered covariates, spatial smoothing using a Gaussian kernel with a full width at half maximum of 8 mm, and spatial normalization to the Montreal Neurological Institute (MNI) space.

### Region of Interest Definition

Following previous network segregation studies (21,24), we defined regions of interest (ROIs) based on the Power-264 brain atlas (41). Each of the 264 ROIs was a 5-mm sphere centered around the atlas coordinates and was assigned to one of 13 distinct brain networks (41). These networks were grouped into three categories: “association” networks (default mode, fronto-parietal, ventral attention, dorsal attention, cingulo-opercular, and salience); “sensorimotor” networks (hand sensory-somatomotor, mouth sensory-somatomotor, visual, and auditory); and “other” networks (subcortical, cerebellar, and memory retrieval), which were not included in analyses (21).

### Network Segregation

For each subject, we calculated the Pearson correlation coefficients between time courses of all ROIs and used Fisher’s *Z*-transformation to convert the values to normally distributed z-scores. Network segregation for each network was defined as the within-network connectivity strength compared to the between-network connectivity strength (21). Specifically, network segregation was the relative difference between the average Fisher’s z-value of connections *within* a given network 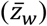 and the average Fisher’s z-value of connections *between* that network and all other networks 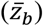 . This difference 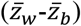 was then normalized by the within-network connectivity 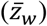

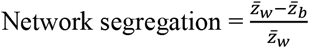

The theoretical maximum of network segregation is 1.0 (ROIs are only connected to ROIs within their network) and 0.0 indicates ROIs are equally connected within and between networks. We calculated the segregation of the sensorimotor and association networks by averaging the network segregation of their constituent networks. Consistent with previous studies, and to avoid potentially spurious negative correlations, only positive connectivity strength was included in our analysis (21,24,27).

### Statistical Analyses

The primary analyses included two hierarchical regressions with either association network segregation or sensorimotor network segregation as the dependent variable. Independent variables in the steps of the regressions were: model 1) mean FD [covariate model to correct for head motion during the fMRI]; model 2) mean FD, age, ASI-drug; and model 3) mean FD, age, ASI-drug, and age × ASI-drug interaction. In follow-up analyses, one of the six association networks were included in turn as the dependent variable. In exploratory analyses in a subsample, we used repeated-measured ANCOVAs to test the effect of XR-NTX treatment on network segregation from baseline to on-treatment (1-2 weeks after first injection) and post-treatment (>1 month after last injection). A *t*-test examined change in ASI-drug from baseline to post-treatment. Exploratory analyses also used partial correlations to examine the relation between baseline network segregation and baseline HAM-D, HAM-A, PANAS subscales, IQ, and WMI.

We conducted statistical tests in SPSS (version 28.0) and used percentile bootstrapping with 5,000 samples. We reported unstandardized betas (*B*)—see tables for standardized betas (β)—and described effect size using proportions of variance explained (*R*^2^) and partial correlations (*r*_partial_). Interactions were probed with simple slopes and Johnson-Neyman regions-of-significance tests (SPSS PROCESS Macro version 4.2). We controlled for Family Wise Error Rate (FWER) or False Discovery Rate (FDR), as appropriate (42). Specifically, we adjusted for having two tests in our main analyses by controlling for FWER with Bonferroni correction (α <

.025). In follow-up analyses, we adjusted for the larger number of tests by controlling for FDR using the Benjamini-Hochberg procedure (42).

## Results

### Association Network Segregation

Model 2 significantly improved upon model 1, *R*^2^_change_ = .19, *p* = .006. ASI-drug was significantly related to lower association network segregation, *B* = -0.26, *p* = .006, 97.5% CI [-0.45, -0.06] (Figure 1). ASI-drug accounted for 17% of the variance in association network segregation and showed a medium-to-large effect, *r*_partial_(36) = -.48. There was a trend of age relating to lower association network segregation, but this did not survive correction for multiple tests, *B* = -0.002, *p* = .046, 97.5% CI [-0.004, 0.0002] (Figure S1 in Supplemental Materials). Model 3 added the ASI-drug x age interaction, but the model fit did not significantly improve, *R*^2^_change_ = .02, *p* = .23. See Table 2 for full results^1^.

**Table 2.**
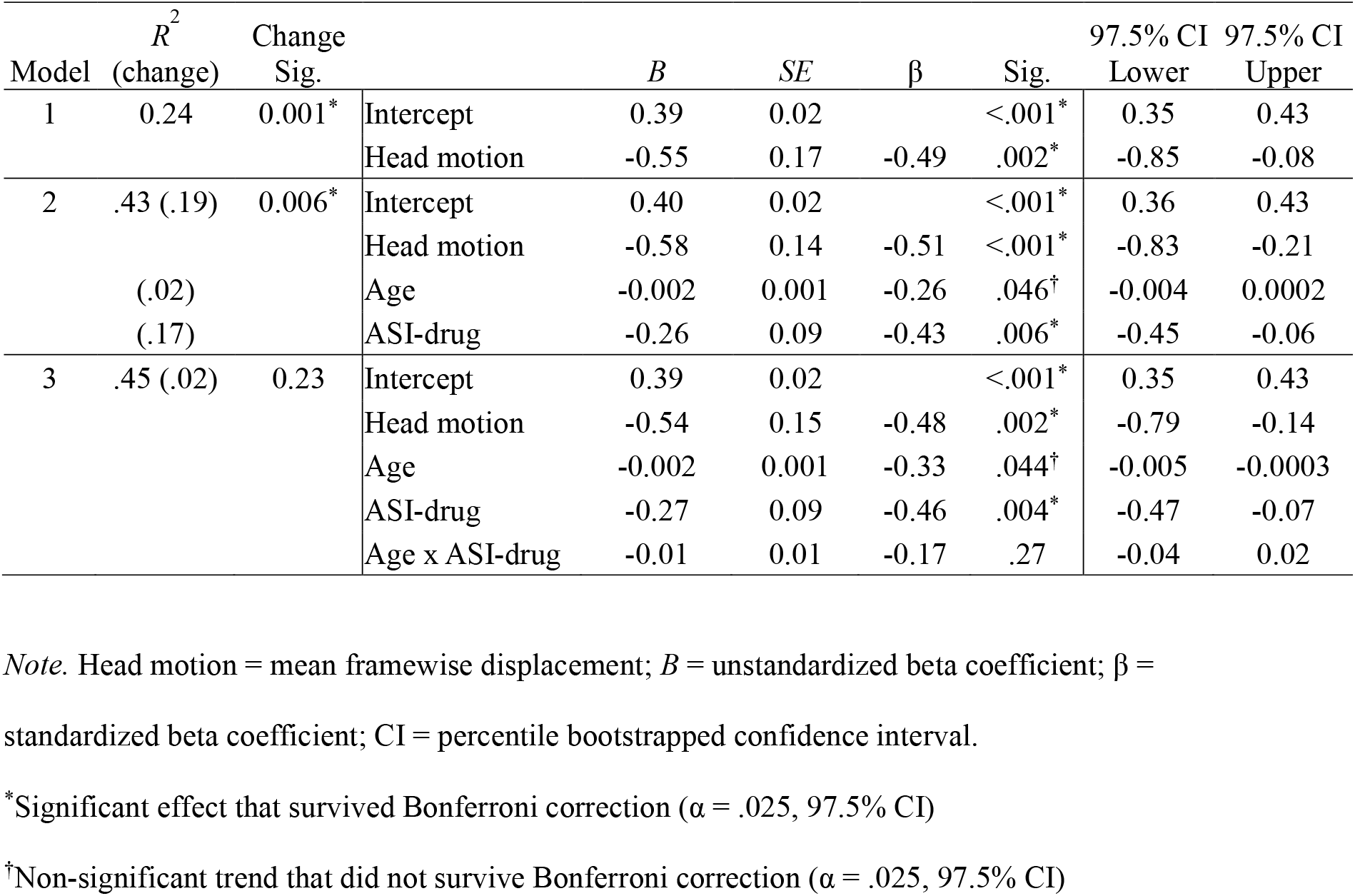
Hierarchical Multiple Regression Predicting Association Network Segregation.

**Figure 1.**
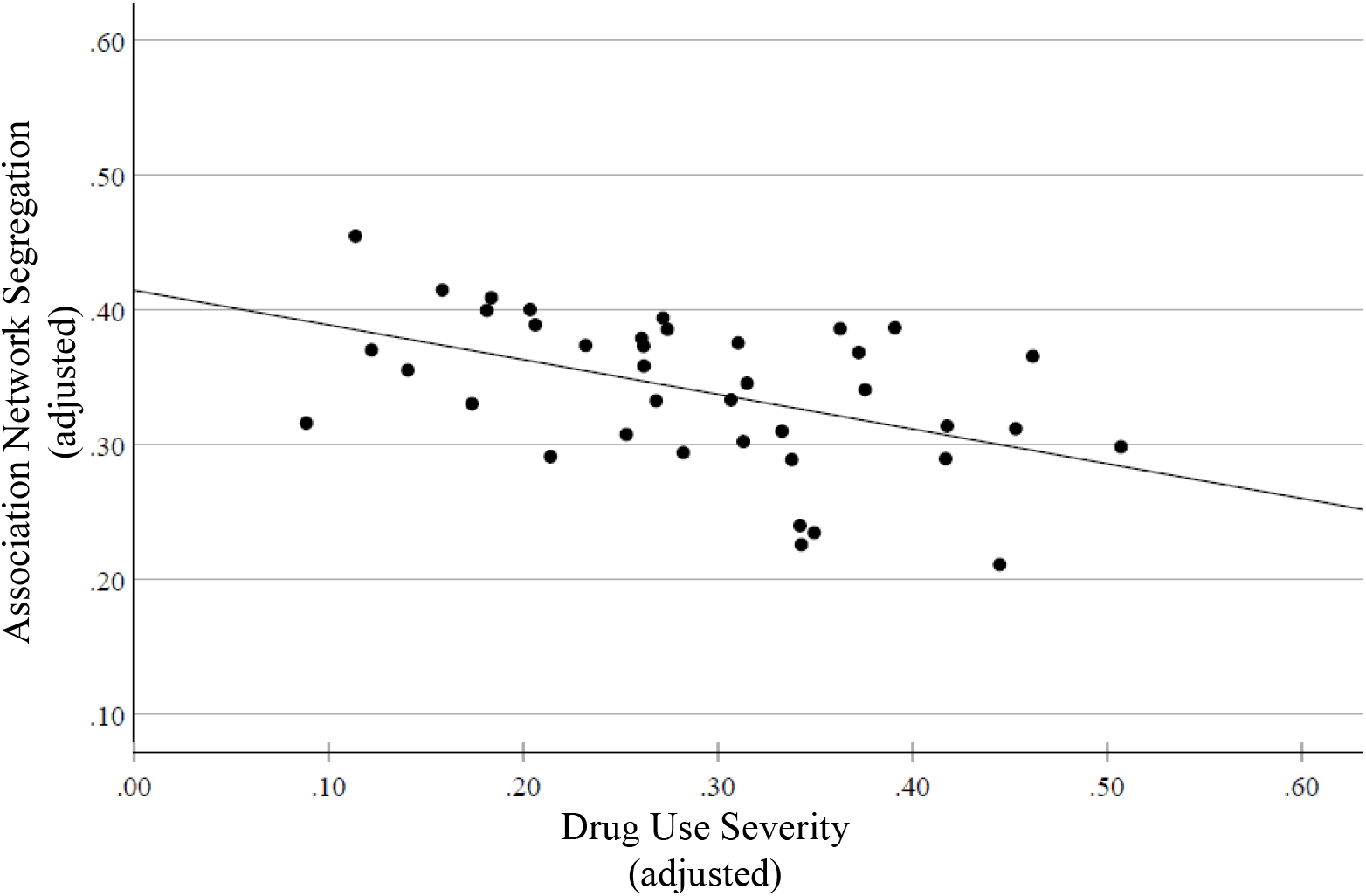
Scatter plot of Association Network Segregation by Drug Use Severity. *Note*. The variables are adjusted by regressing out age and motion (mean framewise displacement). Drug use severity is derived from the Addiction Severity Index’s Drug Composite Score

### Sensorimotor Network Segregation

Model 2 significantly improved upon model 1, *R*^2^_change_ = .20, *p* = .017. Only age was significantly related to sensorimotor network segregation, *B* = -0.004, *p* = .007, 97.5% CI [-0.007, -0.0005], while the effect of ASI-drug was not significant, *B* = -0.09, *p* = .44, 97.5% CI [-0.35, 0.17]. Age accounted for 19% of the variance in sensorimotor network segregation and showed a medium-to-large effect, *r*_partial_(36) = -.45. This effect was qualified by a significant improvement in model fit when adding the ASI-drug x age interaction in model 3, *R*^2^_change_ = .12, *p* = .01. This interaction was on the edge of the corrected significance level, *B* = 0.04, *p* = .026, 97.5% CI [-0.003, 0.079], but we probed it due to the significant improvement in model fit. Tests of simple slopes revealed that ASI-drug was related to lower sensorimotor network segregation at 1 *SD* below mean age, *B* = -0.41, *p* = .019, while there was an opposite but non-significant relation at 1 *SD* above mean age, *B* = 0.33, *p* = .10 (Figure S2). The Johnson-Neyman regions-of-significance tests showed a significant negative effect of ASI-drug at age ≤ 23 (*n* = 14)^2^ and a significant negative effect of age when ASI-drug scores were below the mean (*n* = 18). In the age ≤ 23 group, we found a large negative partial correlation between ASI-drug and sensorimotor network segregation, *r*(10) = -.63, *p* = .016 (Figure 2). Likewise, there was a large negative partial correlation between age and sensorimotor network segregation only in participants with low ASI-drug scores (i.e., below the mean; *n* = 18), *r*_partial_(14) = -.78, *p* < .001 (Figure S3). See Table 3 for full results^3^.

**Table 3.**
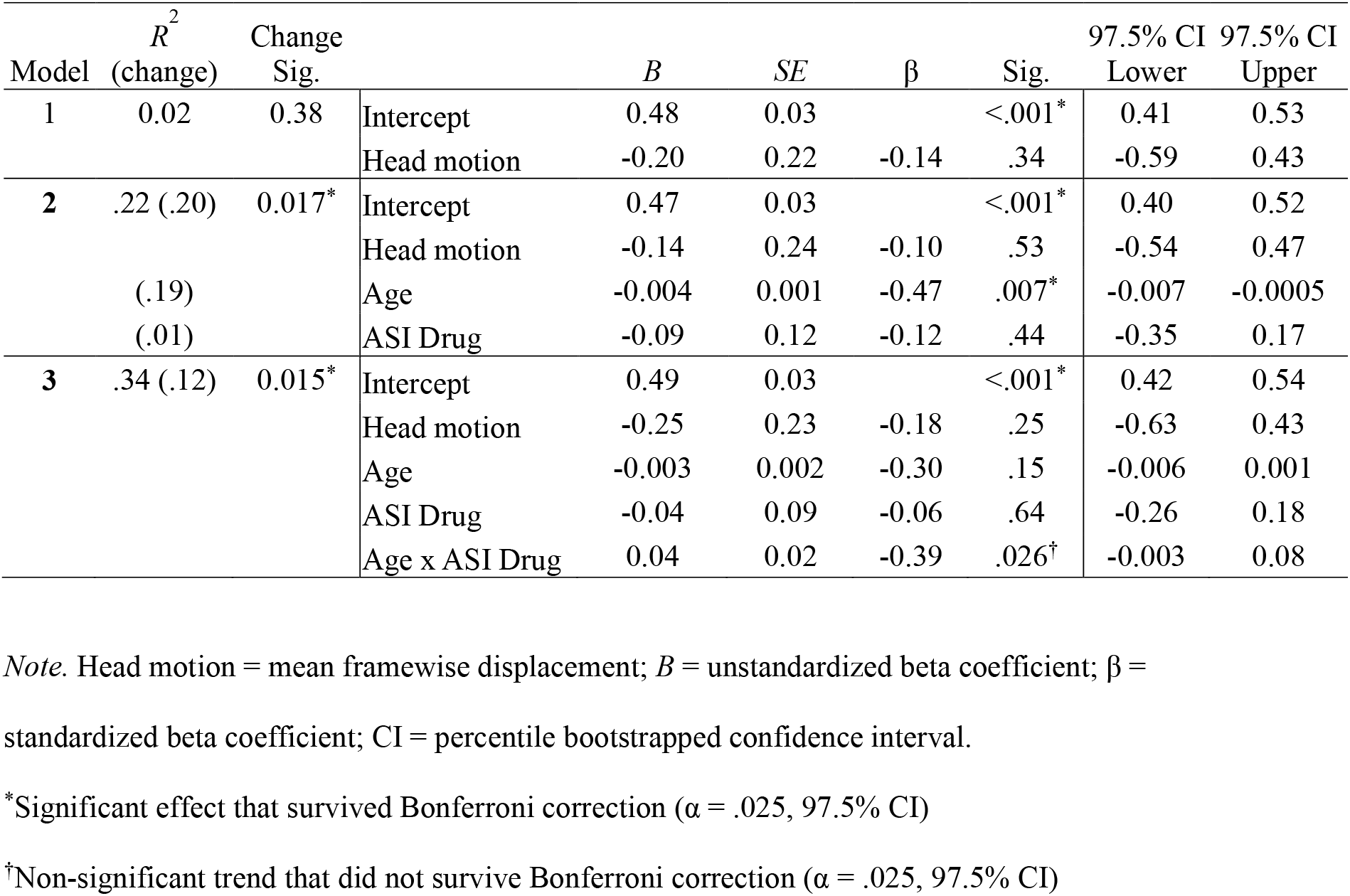
Hierarchical Multiple Regression Predicting Sensorimotor Network Segregation.

**Figure 2.**
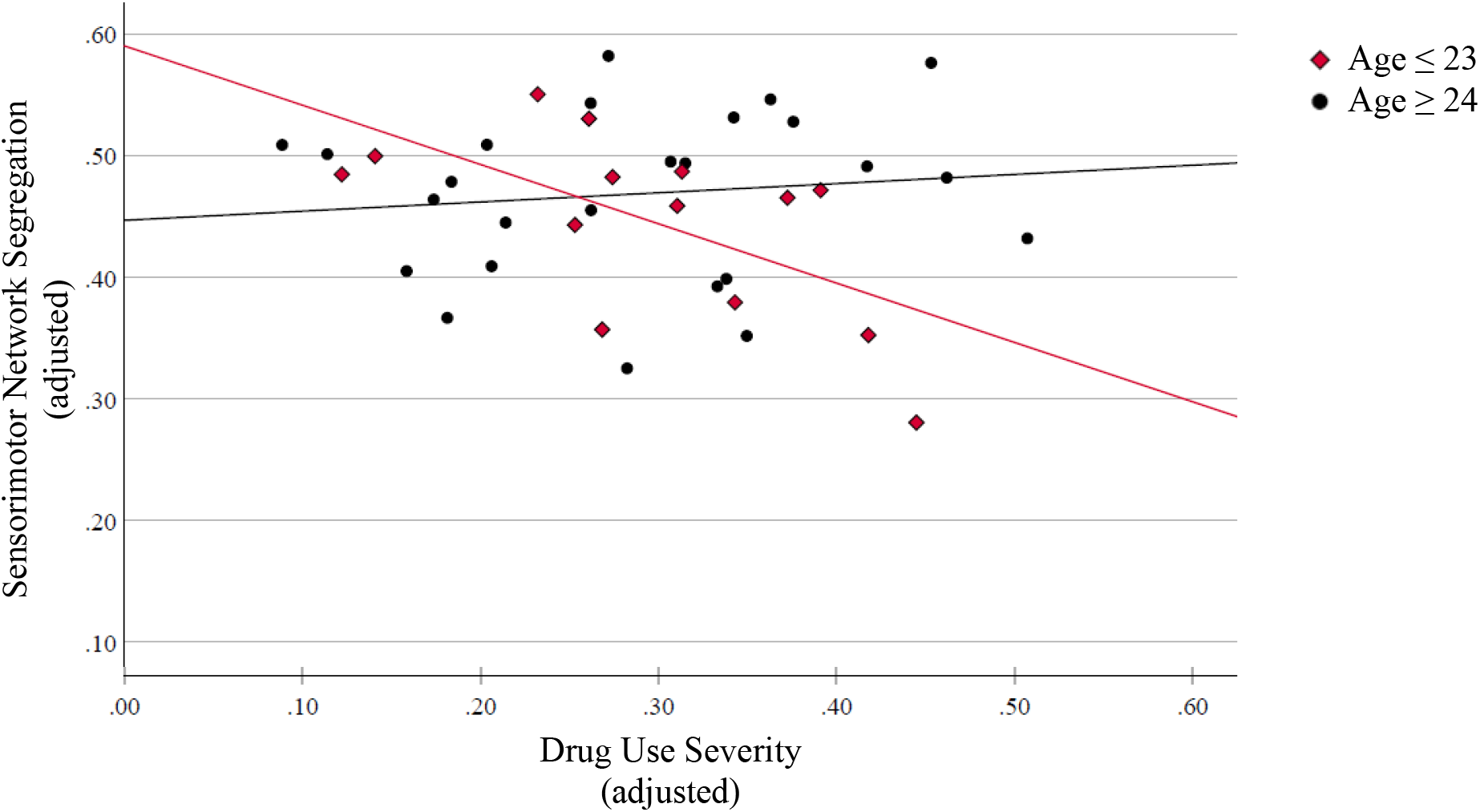
Scatter Plot of Sensorimotor Segregation by Drug Use Severity and Age. Drug Use Severity (adjusted) *Note*. The variables are adjusted by regressing out age and motion (mean framewise displacement). Drug use severity is derived from the Addiction Severity Index’s Drug Composite Score

### Specific Association Networks

Figure 4 shows spring-embedded layouts (Cytoscape version 3.10.1) to visualize the edges and nodes as a function of each specific network.

#### Salience Network

FDR-corrected α = .008. Model 2 significantly improved upon model 1, *R*^2^_change_ = .27, *p* = .002. ASI-drug was significantly related to lower SN segregation, *B* = -0.46, *p* = .004, 97.5% CI [-0.82, -0.16]. ASI-drug accounted for 24% of the variance in SN segregation and showed a large effect *r*_partial_(36) = -.52, *p* < .001 (Figure S4). Age showed a non-significant but trending relation to reduced SN segregation after *p*-value correction, *B* = -0.003, *p* = .040, 97.5% CI [-0.007, 0.000]. In model 3, the ASI-drug x age interaction did not significantly improve model fit, *R*^2^_change_ = .01, *p* = .62.

#### Fronto-parietal Network

FDR-corrected α = .017. Model 2 showed a trend toward improving upon model 1 after correction for multiple tests, *R*^2^_change_ = .18, *p* = .022. There was a significant effect of ASI-drug being related to lower FPN segregation, *B* = -0.39, *p* = .011, 97.5% CI [-0.73, -0.10]. ASI-drug accounted for 18% of the variance in FPN segregation and showed a medium-to-large effect *r*_partial_(36) = -.43, *p* = .005 (Figure S4). Age was not significantly related to FPN segregation, *B* = -0.001, *p* = .688, 97.5% CI [-0.004, 0.003]. In model 3, the ASI-drug x age interaction did not significantly improve model fit, *R*^2^_change_ = .01, *p* = .64.

#### Other Association Networks

None of the other four association networks showed significant effects of ASI-drug on network segregation. See Supplemental Materials for full results from these networks.

### Exploratory Results

#### Network Segregation and Treatment Outcome

Outcome analyses included only participants who completed the post-treatment assessments (*n* = 16). ASI-drug scores significantly reduced from baseline (T1) to post-treatment (T3), *t*(15) = 6.43, *p* < .001, 95% CI [0.14, 0.26], Hedges’ *g* = -1.57. This indicated a large effect of naltrexone treatment, such that 100% of the ASI-drug scores at T3 were below the T1 mean score. In contrast, network segregation for both the association and sensorimotor networks was stable across all three time points (*p*’s > .23; see Supplemental Materials). See Table 4 for means and effect sizes.

**Table 4.** Comparing Mean Network Segregation and Drug Use Severity Across Time.

|  | Time 1 | Time 2 | Time 3 | Time 1<br>vs. Time 2 |  | Time 1<br>vs. Time 3 |  | Time 2<br>vs. Time 3 |  |
| --- | --- | --- | --- | --- | --- | --- | --- | --- | --- |
|  | Mean (*SD*) |  |  | *p* | *g* | *p* | *g* | *p* | *g* |
| Days since Time 1 | -- | 13.0 (7.7) | 112.2 (21.4) | -- |  | -- |  | -- |  |
| Association<br>network segregation | .355 (.066) | .334 (.086) | .332 (.072) | .24 | -0.30 | .16 | -0.36 | .93 | -0.02 |
| Sensorimotor<br>network segregation | .436 (.090) | .428 (.111) | .448 (.071) | .70 | -0.10 | .58 | -0.14 | .43 | 0.20 |
| Drug use severity | .262 (.103) | -- | .062 (.072) | -- |  | <.001 | 1.57 | -- |  |

*Note:* Drug use severity was not collected at Time 2. Time 1 = baseline/pre-treatment, Time 2 = on-treatment (week 2 or 3), Time 3 = post-treatment (>1 month post-injection); *SD* = standard deviation.

#### Network Segregation and Mental Health

We explored associations between mental health variables and networks that related to drug use severity in the previous analyses. Pearson partial correlations (controlling for mean FD, age, ASI-drug) tested whether segregation of the association networks, sensorimotor networks, SN, and FPN were related to depression severity (*n* = 39), anxiety severity (*n* = 38), positive affect (*n* = 24), and negative affect (*n* = 24)^4^. Results showed mostly small and non-significant correlations (Table S1). However, SN segregation significantly and positively correlated with positive affect, *r*(19) = .62, *p* = .0029 (multiple correlation FDR-corrected α = .0031; Figure 3).

**Figure 3.**
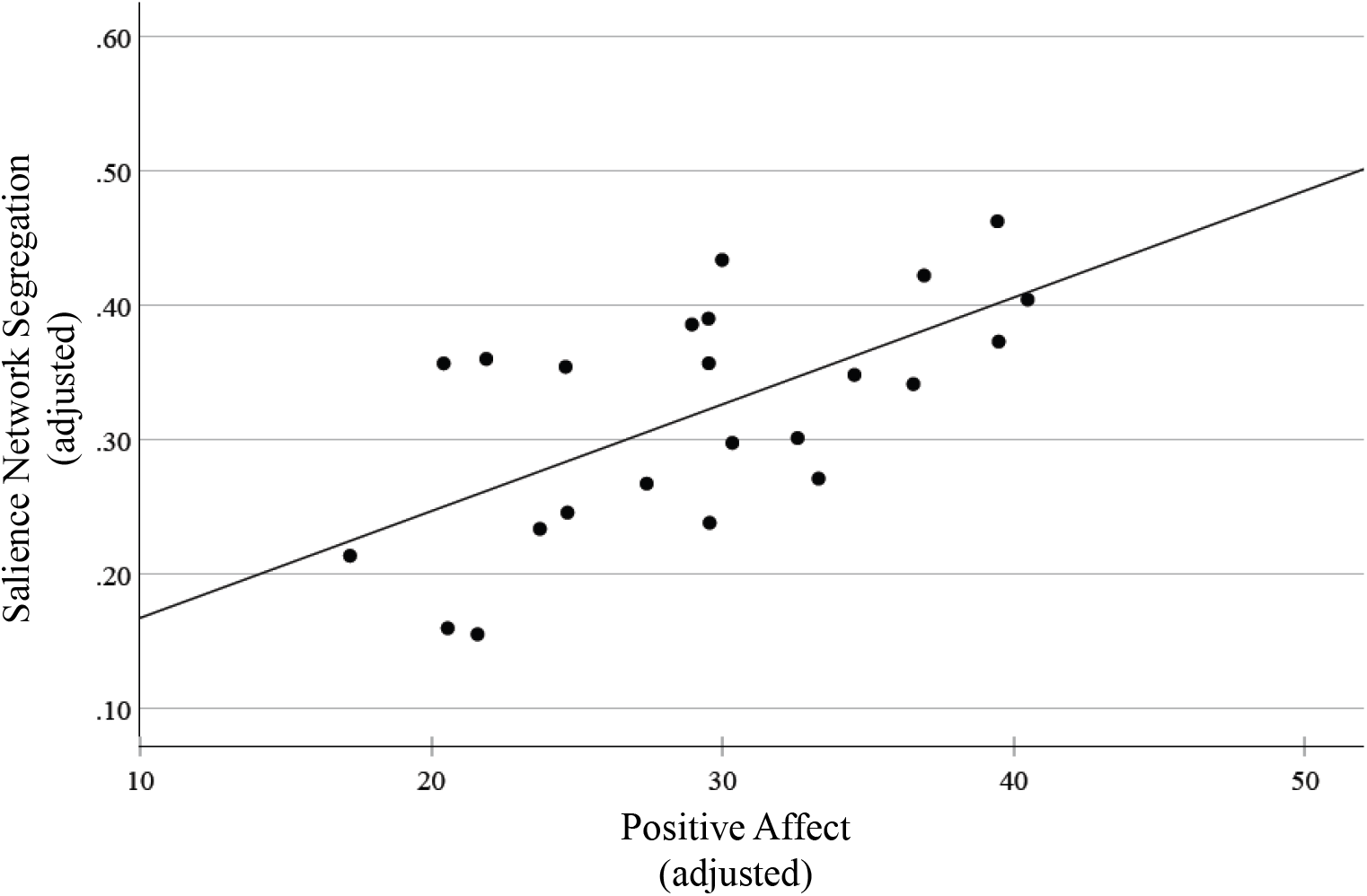
Scatter Plot of Salience Network Segregation by Positive Affect. *Note*. The variables are adjusted by regressing out age and motion (mean framewise displacement). Positive affect = Positive affect subscale of the Positive Affect Negative Affect Schedule.

**Figure 4.**
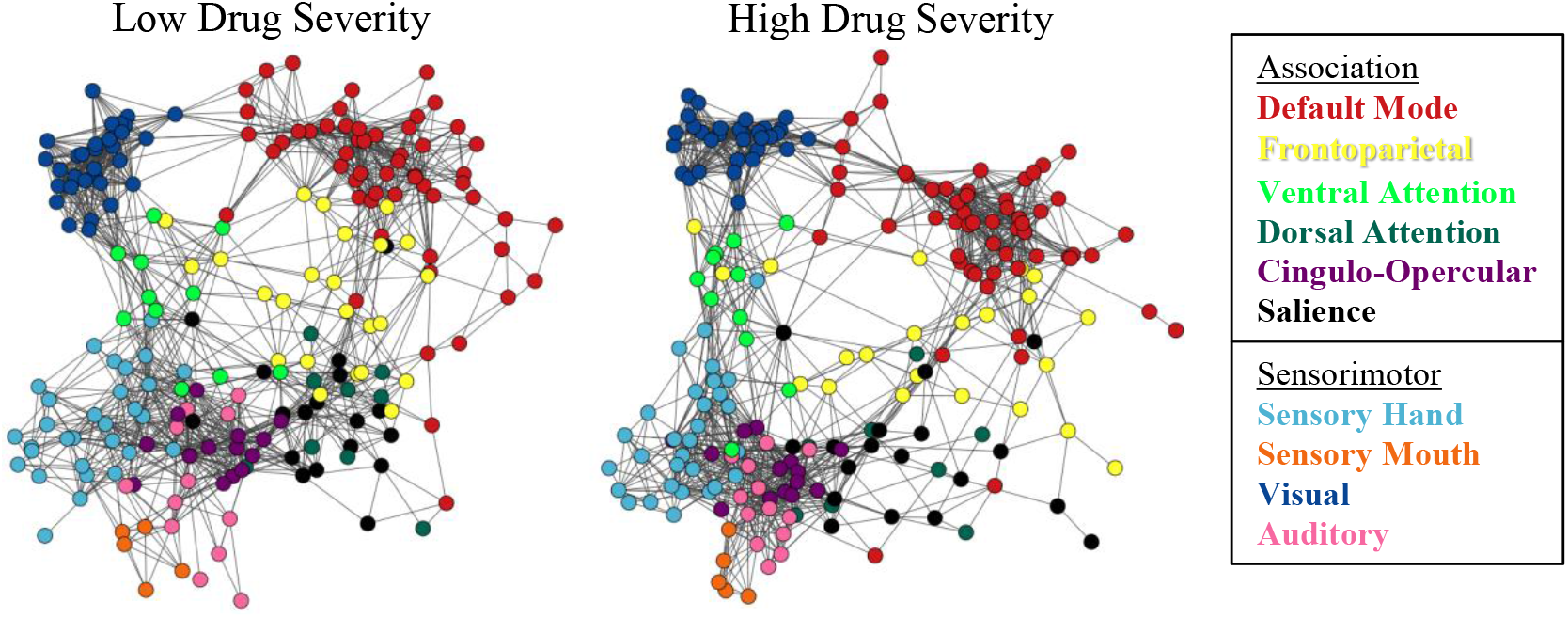
Spring-embedded Layouts of Resting State Functional Connectivity by Network. *Note*. Plots were created using 5% edge threshold. Observe the greater spread between nodes among the Frontoparietal and Salience networks in the high drug severity group, as was substantiated by the statistical results. Low drug severity: Addiction Severity Index Drug Composite ≤ -0.5 *SD* below the mean. High drug severity: Addiction Severity Index Drug Composite ≥ 0.5 *SD* above the mean.

#### Network Segregation and Cognition

Pearson partial correlations (controlling for mean FD, age, ASI-drug) tested whether segregation of the association networks, sensorimotor networks, SN, and FPN were related to IQ (*n* = 40) and working memory (*n* = 26)^5^. Results showed no significant correlations between network segregation and these cognitive measures (Table S1).

## Discussion

This secondary analysis examined drug use severity and brain network segregation in individuals with OUD seeking treatment with XR-NTX. Consistent with our hypothesis, drug use severity was related to lower network segregation in association networks. This was particularly driven by large effects in the FPN and SN, but not in the DMN or other association networks. This highlights the specific vulnerability of certain higher-order cognitive networks to the effects of drug use severity. We observed a trend for older age predicting lower segregation in association networks, but this effect did not reach significance. In contrast, drug use severity was related to lower sensorimotor network segregation in only younger adults. Exploratory analyses revealed a large association between positive affect and greater SN segregation, suggesting a potential link between network organization and emotional well-being. We found no relation between brain network segregation and cognition and no change in segregation during XR-NTX treatment.

In the association networks, we found relations between drug use severity and lower brain network segregation in the FPN and SN, which are often disrupted in OUD. Interestingly, DMN segregation was not associated with drug use severity. In contrast to the task-negative DMN, the task-positive FPN activates during tasks to support cognitive functions such as decision-making, attentional control, and working memory (43). The SN is a dynamic switch between the FPN and DMN, and serves to identify contextually relevant stimuli and integrate it to guide behavior (43–45). The relation of drug use severity to lower network segregation in these networks implies potential consequences for cognitive performance and control over highly salient stimuli. Indeed, individuals with OUD have impaired executive functioning (46), which may be partially explained by reduced FPN segregation. As we did not find an association between FPN network segregation and working memory, future studies should examine other executive functions. Alternatively, it is possible that OUD-related damage decoupled FPN segregation from executive function, as suggested by a study showing that association network segregation positively correlated with working memory in healthy control individuals but not in people with alcohol use disorder (28). Future research should investigate if changes in network segregation can act as a biomarker for cognitive impairment in OUD and whether targeting network organizations could improve these deficits. Longitudinal studies are needed to see if reduced FPN and SN segregation predicts worsening cognitive function or if treatment targeting network organizations could improve these deficits.

The link between sensorimotor network segregation and drug use severity was more nuanced. Although there was no general association, we found that higher drug use severity was related to lower sensorimotor network segregation in the younger participants (≤23 years old). This may suggest that the brains of younger adults with OUD are more vulnerable to drug-related changes in sensorimotor network segregation. Alternatively, underlying differences in sensorimotor network segregation may influence drug use behavior in young adults—who have yet to reach full maturity in prefrontal control of behavior (47). Prior cross-sectional research suggests that brain network segregation in healthy adults declines with age (21). Our work shows that the age-related decrease in sensorimotor network segregation is decoupled in people with OUD who have higher drug use severity. This was due to lower segregation among the younger participants with more severe use. Our results align with previous research indicating that opioid use and craving are associated with less sensorimotor within-network connectivity (48,49). Possible mechanisms for the relation between sensorimotor network connectivity and opioid use severity include *dys*connectivity in these networks contributing to cognitive dysfunction and impulsivity (50). Further, abnormalities in networks that govern motor and sensory aspects of drug use behaviors may increase the automaticity and motivational pull of such behaviors (51). Although our results suggest that age and severe drug use problems impact the topography of sensorimotor networks, larger and longitudinal samples are needed to confirm developmental differences.

Exploratory analyses showed that higher network segregation of the SN was associated with greater self-reported positive affect. Given the role of the SN in detecting and integrating rewarding, engaging, and emotional stimuli (52), a relatively segregated and efficient SN may directly support positive mood states. Individuals with a more optimal balance of within versus between network SN connectivity may more easily detect positive stimuli in the environment and more fully engage with positive emotions internally. Further, greater SN segregation may support emotion regulation skills (53,54), which increases positive affect (55). One study of healthy young adults found a link between positive affect and *lower* SN within-network rsFC (56), which affirms the association although in the opposite direction. However, network segregation is the *balance* of within versus between network connectivity, and people with OUD may have different neural underpinnings of positive affect. SN segregation was not associated with depression, anxiety, or negative affect. SN’s relation to positive affect but not negative affect is consistent with other work showing differential networks involved in positive and negative affect (56,57). The dissociation also aligns with the expected small (and non-significant) correlation between positive and negative affect in our sample and others (36,58).

In this first study to test network segregation response to treatment, we did not observe a significant change across three months of XR-NTX treatment. This indicates some short-term stability of network segregation in treatment-seeking people with OUD, despite a significant decrease in recent drug use severity. Network segregation may not be as malleable to XR-NTX as self-reported drug severity, which encompasses behavior and subjective ratings. Although XR-NTX alters functional brain response (e.g., reduced cue-reactivity) (32), other interventions may be more suitable for altering connectivity within and between brain networks. Non-invasive neurostimulation is one promising tool. For example, transcranial direct stimulation in people with methamphetamine use disorder modulated rsFC of the DMN, FPN, and SN, which was associated with reduced methamphetamine craving (59). Research suggests that stimulation deeper in the brain through low intensity focused ultrasound may alter network rsFC of subcortical regions involved in SUD (60,61). Further, the psychedelic substance psilocybin may open a window for flexibility in network connectivity such as in decreased network modularity associated with reduced depression (62). Future work could examine whether psilocybin-induced flexibility enables a complementary normalization of network segregation/integration in people with SUDs.

This study comes with limitations commonly encountered with secondary analyses that limit the generalizability of the results. The study sample was relatively small and from predominantly White and male individuals with heroin and prescription opioid use, reflecting the demographics and drug of choice from the period that preceded the broad introduction of synthetic opioids to the illicit drug supply in the United States (63,64). We lacked a healthy control sample and could not directly establish whether OUD generally relates to brain network segregation deficits. Nevertheless, the mean brain network segregation measures in our OUD sample were significantly lower than the healthy control and alcohol use disorder groups in our previous study (28) (*p*s < .001; see Supplemental Results for ANCOVAs and marginal means). However, it is difficult to directly compare network segregation across these studies because the current study used a shorter resting-state fMRI (∼5 minutes) compared to the previous study (∼15 minutes) (65). The associations between brain network segregation and drug use were cross-sectional, and causality and its direction are unknown. Although we saw no change in network segregation during the study period, datasets that measure rsFC prior to and during substance use will better describe changes in network segregation.

## Conclusion

In the first study of brain network segregation in people with OUD, we showed that those with high recent drug use severity had lower network segregation in association networks, especially the FPN and SN. Inefficient executive functioning and salience processing may contribute to or be impacted by ongoing substance use problems in people with OUD. Sensorimotor network segregation was only related to drug use severity in young adults, indicating a potential vulnerability in these individuals. Our findings provide better understanding of reduced efficiency and premature aging of brain networks in OUD and are a step towards developing interventions to address the impact of OUD on the brain network architecture.

## Footnotes

1 Adding IQ and ASI alcohol severity as covariates showed nearly identical results as the main analyses and no change in their interpretation. Neither IQ (*B* = 0.0003, *p* = .67, 97.5% CI [-0.002, 0.001]) nor alcohol severity (*B* = 0.23, *p* = .09, 97.5% CI [-0.15, 0.59]) were significantly related to association network segregation.

2 There was also a significant positive effect of ASI-drug on sensorimotor network segregation at age > 44, but this effect was less interpretable because this group included only *n* = 4 participants.

3 Adding IQ and ASI alcohol severity as covariates showed nearly identical results as the main analyses and no change in their interpretation. Neither IQ (*B* = 0.001, *p* = .50, 97.5% CI [-0.002, 0.003]) nor alcohol severity (*B* = 0.17, *p* = .41, 97.5% CI [-0.35, 0.78]) were significantly related to sensorimotor network segregation.

4 Sample sizes varied based on available data. Depression and anxiety scores were missing for 1 and 2 participants, respectively. The PANAS (positive and negative affect) was only collected for the first half of the study.

5 Sample sizes varied based on available data. The working memory was only collected for the first half of the study.

## Notes

Funding: This work was supported by NIH/NIDA T32DA028874 (Hager); NIH/NIA T32AG076411 (Ramos-Rolón); Commonwealth of Pennsylvania C.U.R.E. Addiction Center of Excellence: Brain Mechanisms of Relapse and Recovery (Childress); NIH AA031088, NIH AA031337, and NIH DA046345-05W1 (Wiers); NIH/NIDA K01DA051709, Brain & Behavior Research Foundation NARSAD Young Investigator Grant #30780 (Shi);

### Competing Interest Statement

The authors have declared no competing interest.

