## Supplemental Materials for "Brain Network Segregation is Associated with Drug Use Severity in Individuals with Opioid Use Disorder"

### Supplemental Method

#### *Study Procedures*

Opioid detoxification was conducted under the care of the study physician and confirmed via an opioid-negative urine drug screen and naloxone challenge to ensure the absence of opioid withdrawal signs. For nearly all participants, the first of three monthly XR-NTX injections was administered immediately after the baseline MRI scan ($n$ = 29) or within one week ($n$ = 5). The remaining participants received their first injection 3-to-5 weeks after the MRI scan ($n$ = 2) or withdrew from the study before injection ($n$ = 4).

#### *Statistical Analyses*

FWER with Bonferroni correction ($\alpha < .025$) was used in the main analyses to kept the risk of a single type I error (false rejection of the null hypothesis) at the overall desired α level ($\alpha = .05$). Because of the larger number of tests in the follow-up analyses, Benjamini-Hochberg procedure was used to apply a different α to each test based on $p$-value rank and kept the proportion of type I error among all significant results at the overall desired α level ($\alpha = .05$), while minimizing type II error (1).

### Supplemental Results

#### *Default Mode Network*

FDR-corrected $\alpha = .025$. Model 2 (ASI-drug and age added) showed a trend toward improving upon the covariate model (motion) after correction for multiple tests, $R^2_{change} = .12$, $p = .05$. Within Model 2, the effect of ASI-drug on network segregation was not significant, $B = -0.19$, $p = .17$, 97.5% CI [-0.51, 0.11]. However, age was significantly related to decreased network segregation, $B = -0.004$, $p = .004$, 97.5% CI [-0.007, -.001], showing a small effect,

$r_{partial}(36) = -.22$. In Model 3, the addition of the ASI-drug x age interaction did not significantly improve model fit, $R^2_{change} = .02$, $p = .29$.

***Ventral Attention Network***

FDR-corrected α = .033. Model 2 did not significantly improve on the covariate model, $R^2_{change} = .08$, $p = .23$. Within model 2, the effect of ASI-drug on network segregation was not significant, *B* = -0.28, *p* = .13, 96.7% CI [-0.64, 0.14] and neither was the effect of age (*B* = 0.002, *p* = .39, 96.7% CI [-0.003, 0.005]). In model 3, the addition of the ASI-drug x age interaction did not significantly improve model fit, $R^2_{change} = .06$, $p = .12$.

***Cingulo-opercular Network***

FDR-corrected α = .042. Model 2 did not significantly improved on the covariate model, $R^2_{change} = .08$, $p = .20$. Within model 2, the effect of ASI-drug on network segregation was not significant, *B* = -0.17, *p* = .31, 95.8% CI [-0.51, 0.13], and neither was the effect of age, *B* = -0.003, *p* = .095, 95.8% CI [-0.006, 0.001]. In model 3, the addition of the ASI-drug x age interaction did not significantly improve model fit, $R^2_{change} = .02$, $p = .31$.

***Dorsal Attention Network***

FDR-corrected α = .05. Model 2 did not significantly improve on the covariate model, $R^2_{change} = .02$, $p = .61$. Within model 2, the effect of ASI-drug on network segregation was not significant, *B* = -0.05, *p* = .76, 95% CI [-0.40, 0.30] and neither was the effect of age, *B* = -0.002, *p* = .24, 95% CI [-0.005, 0.001]. In model 3, the addition of the ASI-drug x age interaction did not significantly improve model fit, $R^2_{change} = .002$, $p = .80$.

***Network Segregation across Treatment***

Specifically, there was no effect of time on association network segregation, $F(2,22) = 1.02$, $p = .37$, $\eta_p^2 = .07$, and no interaction between time and baseline age, $F(2,22) = 1.45$, *p* =

.25, $\eta_p^2$ = .09. Similarly, there was no effect of time on sensorimotor segregation, $F(2,28) = 0.42$, $p = .67$, $\eta_p^2 = 0.03$, and no interaction between time and baseline age, $F(2,22) = 1.56$, $p = .23$, $\eta_p^2 = .10$.

***Network Segregation Group Comparisons***

We compared network segregation in the opioid use disorder (OUD) group to that of our previous study's alcohol use disorder (AUD) and healthy control (HC) groups (2). However, note that this analysis is limited by the difference in scan time of the resting-state functional magnetic resonance imaging between the two studies (~5 minutes for OUD and ~15 minutes for AUD and HC), as longer scan times result in higher network segregation (3). Analysis of covariance (ANCOVA) showed a main effect of group controlling for age and mean framewise displacement (mean FD) to predict segregation from group (HC, AUD, and OUD).

There was a significant effect of group on association network segregation, $F = 20.1$, $p < .001$, partial eta$^2$ = 0.24. Estimated marginal means showed that association network segregation was significantly lower for the OUD group ($M = .31$, $SE = .01$) compared to the AUD group ($M = .36$, $SE = .01$, $p = .008$) and the HC group ($M = .40$, $SE = .01$, $p < .001$), which were also different from each other ($p < .001$). For sensorimotor network segregation, there was a significant effect of group, $F = 35.5$, $p < .001$, partial eta$^2$ = 0.36. Estimated marginal means showed that sensorimotor network segregation was significantly lower for the OUD group ($M = .43$, $SE = .01$) compared to the AUD group ($M = .52$, $SE = .01$, $p < .001$) and the HC group ($M = .57$, $SE = .01$, $p < .001$), which were also different from each other ($p = .002$). See Supplemental Figure S5.

**Supplemental Figures**

**Figure S1**

*Scatter Plot of Association Network Segregation by Age*

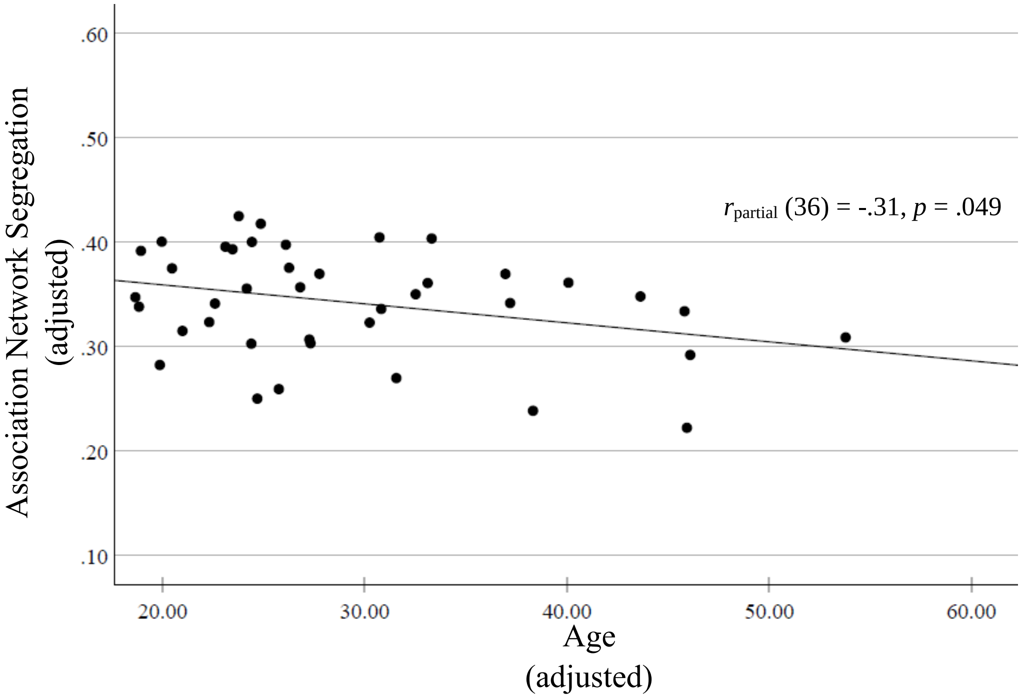


*Note.* The variables are adjusted by regressing out drug use severity (Addiction Severity Index) and motion (mean framewise displacement).

**Figure S2**

*Interaction Plot of Sensorimotor Network Segregation by Drug Use Severity and Age*

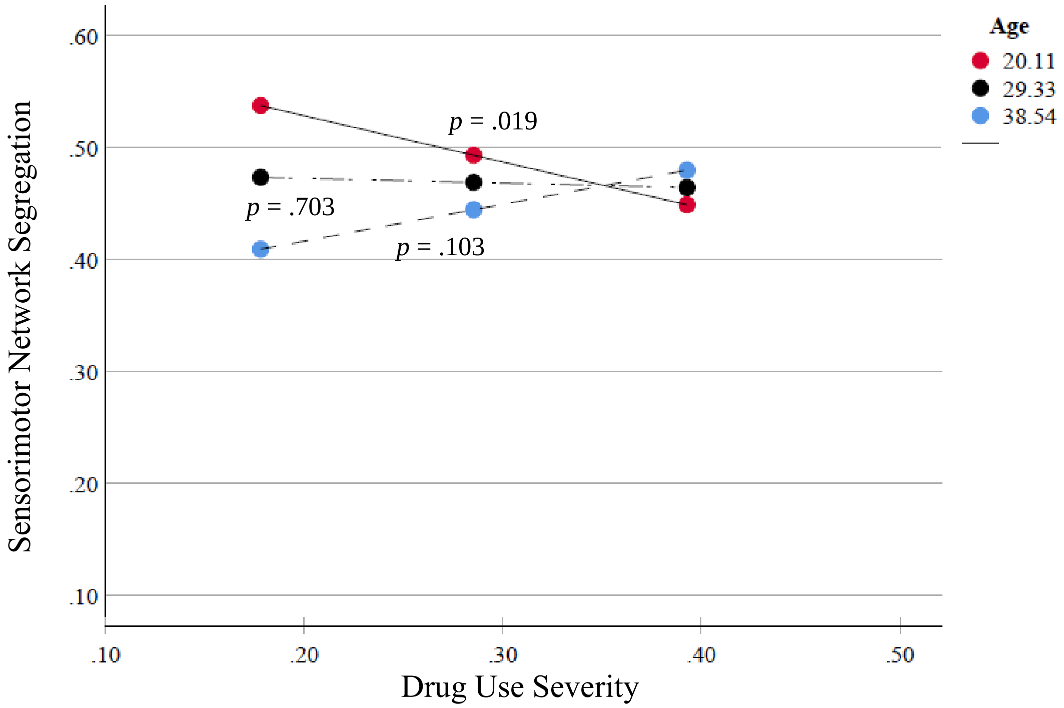


*Note.* Simple slopes are shown here at mean age (29.33 years), 1 standard deviation below the mean (20.11 years), and 1 standard deviation above the mean (38.54 years).

**Figure S3**

*Scatter Plot of Sensorimotor Network Segregation by Age and Drug Use Severity*

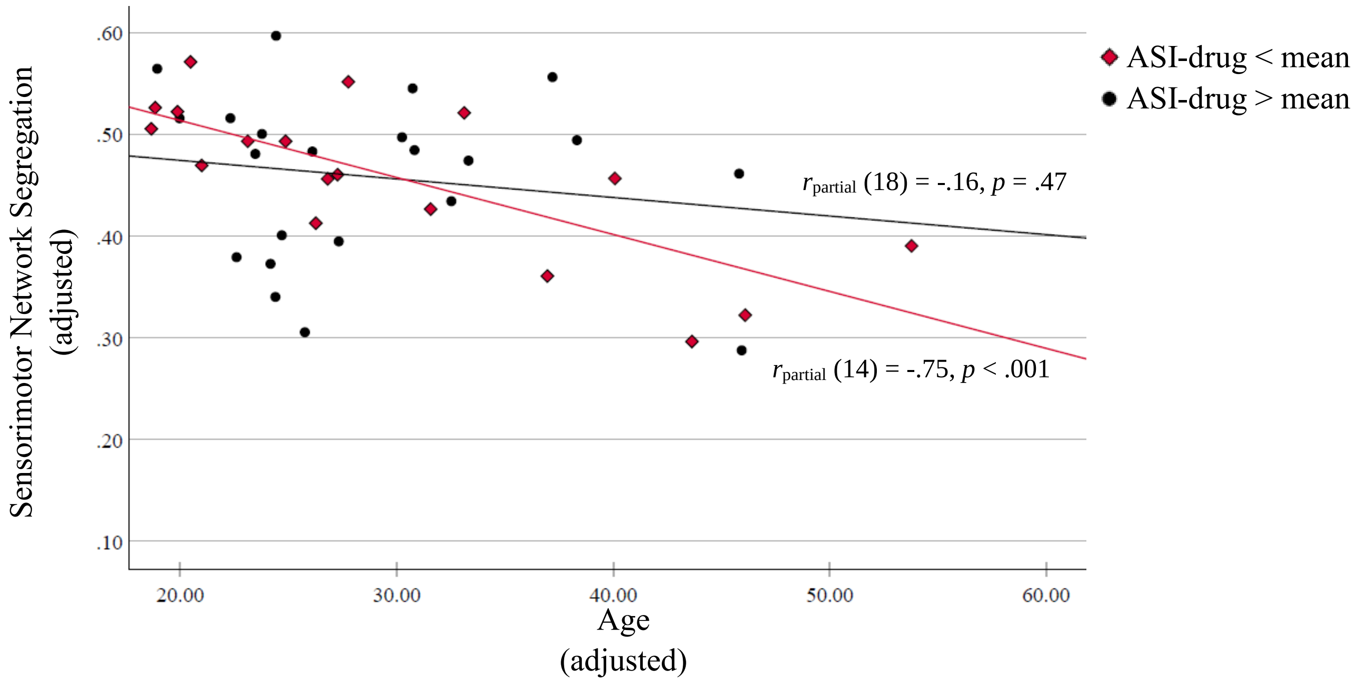


*Note.* ASI-drug = Addiction Severity Index Drug Composite. The variables are adjusted by regressing out drug use severity (ASI-drug) and motion (mean framewise displacement).

**Figure S4**

*Scatter Plots of Selected Networks by Drug Use Severity*

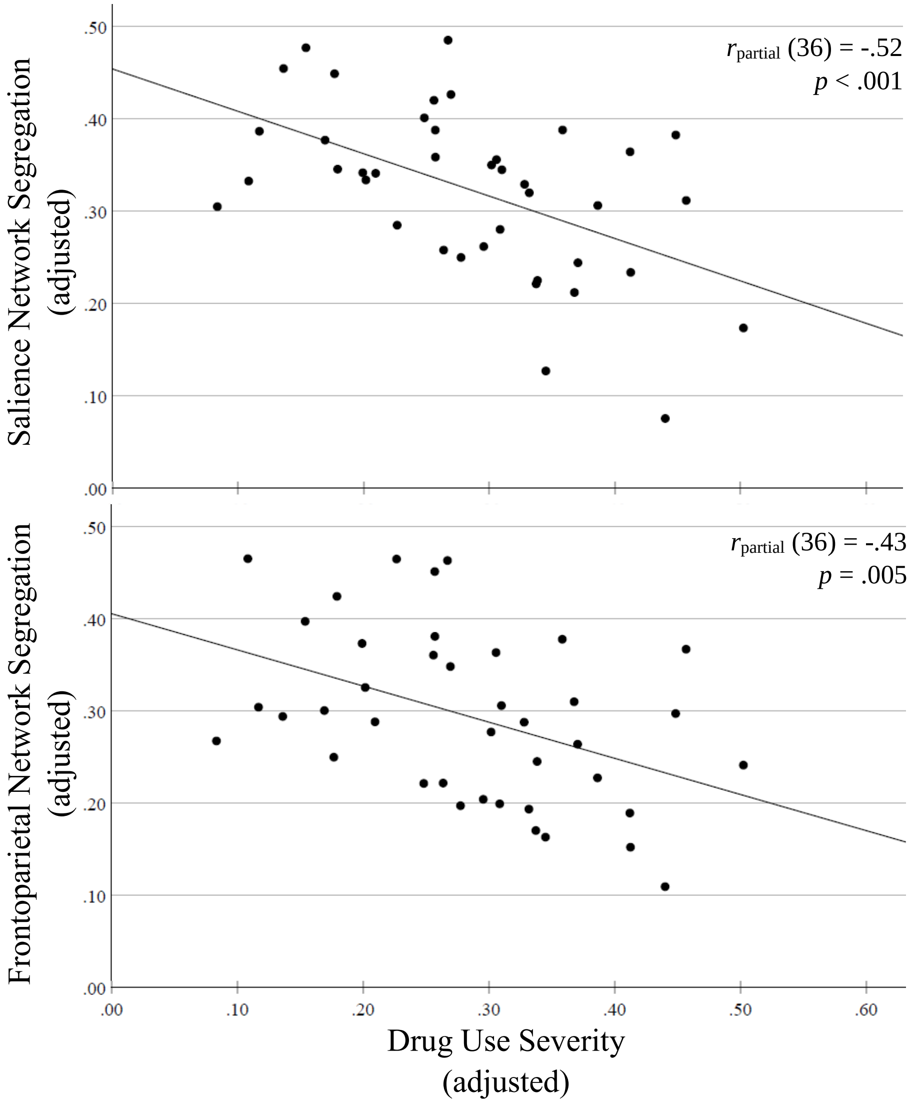


*Note.* The variables are adjusted by regressing out age and motion (mean framewise displacement).

**Figure S5**

*Estimated Marginal Means of Association and Sensorimotor Network Segregation by Group*

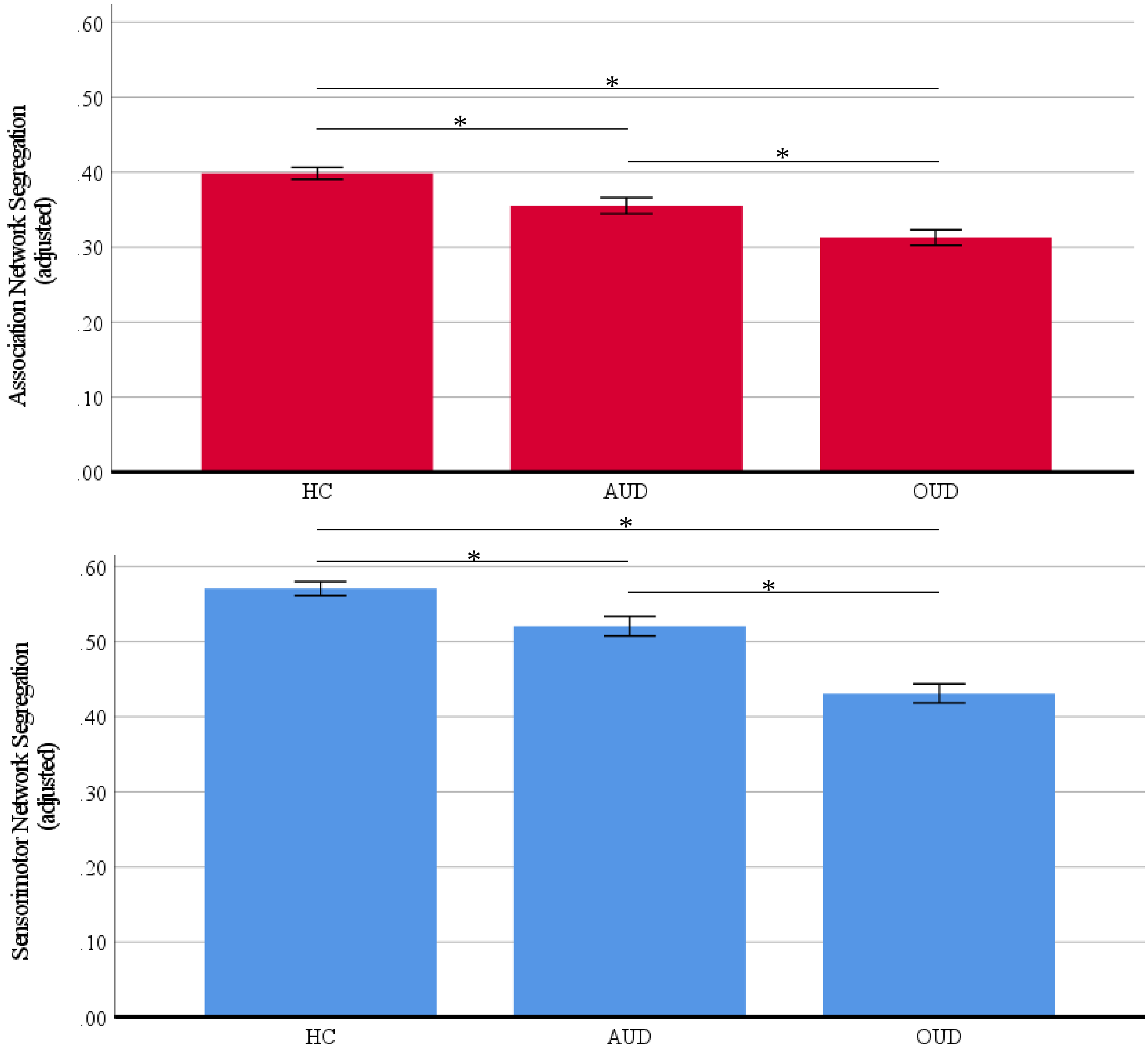


$^{*}p < .01$

$^{**}p <$ 001

*Note.* Network segregation estimated marginal means are adjusted for age and motion (mean framewise displacement). Error bars represent ± 1 standard error of the mean.

**Table S1**

*Partial Correlations of Network Segregation with Mental Health and Cognitive Variables*

| | | Association network segregation | | Sensorimotor network segregation | | Salience network segregation | | Frontoparietal network segregation | |
|---|---|---|---|---|---|---|---|---|---|
| | *n* | *r* | *p* | *r* | *p* | *r* | *p* | *r* | *p* |
| Depression | 39 | -.18 | .30 | -.12 | .49 | -.03 | .88 | -.13 | .46 |
| Anxiety | 38 | -.20 | .24 | -.13 | .45 | -.04 | .84 | -.17 | .32 |
| Positive affect | 24 | .31 | .18 | .31 | .17 | .62 | .003* | .03 | .89 |
| Negative affect | 24 | .16 | .48 | -.05 | .83 | .02 | .93 | -.08 | .74 |
| IQ | 26 | -.06 | .72 | .09 | .59 | -.18 | .29 | -.16 | .34 |
| Working memory | 26 | .10 | .64 | .07 | .74 | -.13 | .54 | .08 | .73 |

*Note.* Partial correlations adjust for age, Addiction Severity Index – Drug Composite score, and mean framewise displacement (head motion). Depression = Hamilton Depression Rating Scale. Anxiety = Hamilton Anxiety Rating Scale. Positive affect = subscale of the Positive Affect Negative Affect Scale. Negative affect = subscale of the Positive Affect Negative Affect Scale. IQ = Intelligence quotient. Working memory = Working Memory Index.

$^{*}p < .01$
